# Engineering a High-Fidelity CRISPR-Cas13d System for Targeted RNA Depletion in *Plasmodium falciparum*

**DOI:** 10.64898/2026.11.09.765308

**Authors:** Rafael Serrano-Durán, Diana López-Farfán, Miguel A. Moreno-Mateos, Elena Gómez-Díaz

**Affiliations:** Instituto de Parasitología y Biomedicina López-Neyra, Consejo Superior de Investigaciones Científicas (IPBLN, CSIC), Granada, Spain; Centro Andaluz de Biología del Desarrollo, Pablo de Olavide University/CSIC/Junta de Andalucía (CABD, UPO-CSIC-JA), Sevilla, Spain

**Keywords:** Human malaria, Functional Genomics, Genome editing, Gene Knockdown

## Abstract

Functional genomics in *Plasmodium falciparum* is severely constrained by the evolutionary absence of an endogenous RNA interference pathway and the requirement for time-consuming direct target locus modification by conventional conditional genetic engineering systems. To overcome these limitations, we established a programmable post-transcriptional gene silencing system in *P. falciparum* using High-Fidelity RfxCas13d (HF-RfxCas13d), an RNA-guided RNA nuclease that has been previously tested in several eukaryotic hosts. Using CRISPR-SpyCas9 we integrated a HF-RfxCas13d expression cassette at the dispensable *pfs47* locus and generated a stable marker-free transgenic line capable of constitutive HF-RfxCas13d cytosolic expression without impairing intraerythrocytic asexual growth. Episomal delivery of single or dual guide RNAs (gRNAs) achieved significant target mRNA depletion (41%-76%) across three endogenous loci *atp4*, *ap2-exp*, *ap2-g* genes. Applying this system to the master transcriptional regulator of gametocytogenesis (*ap2-g*) resulted in transcript depletion that functionally impaired sexual differentiation in committed parasites, reducing significantly the Sexual Conversion Rate a 70%. Overall, our novel HF-Cas13d line provides a flexible, scalable, and non-locus-modifying addition to the *P. falciparum* genetic toolbox to accelerate functional genomics and malaria drug target identification.

## 1. Introduction

Malaria remains a critical global public health challenge, responsible for an estimated 282 million clinical cases and 610,000 deaths in 2024 [1]. The overwhelming majority of severe disease and mortality is caused by *Plasmodium falciparum*. Despite extensive control initiatives, global progress toward the World Health Organization Global Technical Strategy for Malaria (GTS) 2016-2030 targets has stalled, exacerbated by extreme weather events, socio-political instability, and operational disruptions. Current interventions are further threatened by widespread vector resistance to pyrethroid insecticides and the emergence of parasite partial resistance to artemisinin-based combination therapies (ACTs) [1]. Reversing this stagnation requires scalable, programmable genetic tools to interrogate basic parasite biology and accelerate the identification and validation of novel therapeutic targets.

Functional genomics in *P. falciparum* has historically been hindered by the parasite’s atypical biological architecture, including an extremely AT-rich genome (>80.6%), a low proportion of introns, and the absence of canonical non-homologous end joining (NHEJ) machinery [2, 3]. High-throughput piggyBac transposon insertional mutagenesis has revealed that almost half of protein-coding genes in *P. falciparum* are non-mutable and essential for asexual blood-stage proliferation [4]. Because the parasite is haploid during its intraerythrocytic development, direct gene knockouts of essential loci are lethal, preventing the isolation of viable mutant lines for phenotypic characterization for half of the parasite genes. While CRISPR-Cas9 technology has substantially improved genetic editing in *P. falciparum*, investigating essential genes still relies on conditional systems acting at the DNA, RNA, or protein levels [5]. Furthermore, the lack of an endogenous RNA interference (RNAi) pathway (owing to the evolutionary absence of Dicer and Argonaute 2 (Ago2) orthologs) prevents the use of standard small interfering or short hairpin RNAs for direct post-transcriptional gene silencing [6,7]. Although an engineered, Ago2 system has successfully enabled gene silencing in the murine malaria parasite *P. berghei*, this approach has not yet been tested in *P. falciparum* [8]. To bypass this limitation, knockdown and conditional platforms such as destabilizing domains (DD) [9], TetR-aptamer modules (TetR-DOZI) [10], *glmS* ribozymes [11], auxin-inducible degron (AID) [12], knock sideways (KS) [13] and dimerizable Cre recombinase (DiCre) [14] have been adapted to *P. falciparum*. However, these approaches present notable limitations: transcript- or protein-targeted conditional systems require time-consuming locus modification for every target gene. Besides, they often exhibit variable knockdown efficiency, whereas DiCre-mediated gene excision requires complex genomic engineering to introduce *loxP* sites flanking target sequences [5, 15, 16]. Consequently, a major technological gap persists for a programmable knockdown platform that enables flexible functional interrogation of endogenous genes without requiring direct genetic modification of target loci.

Members of the Class 2 Type VI CRISPR-Cas superfamily, uniquely characterized by two Higher Eukaryotes and Prokaryotes Nucleotide-binding domain (HEPN) endoribonuclease domains, act as programmable, RNA-guided RNA-targeting nucleases [17, 18]. Among these, RfxCas13d (derived from *Ruminococcus flavefaciens* strain XPD3002) has emerged as an exceptionally compact (∼930 amino acids) and efficient tool for transcriptome engineering in diverse eukaryotic models. This tool operates independently of protospacer adjacent motif (PAM) or protospacer flanking sequence (PFS) restrictions [17–21]. Upon target RNA binding, RfxCas13d undergoes a conformational change that brings its two catalytic HEPN domains together, inducing precise *cis*-cleavage of the target transcript. However, target recognition simultaneously converts wild-type Cas13 orthologs into activated, non-specific endoribonucleases capable of cleaving non-targeted bystander transcripts in *trans* [17, 18, 22, 23]. This nonspecific "collateral" activity has been associated with non-specific ribosomal RNA (rRNA) degradation, off-target transcriptome disruption, growth retardation, and severe *in vivo* toxicity in several eukaryotic hosts, such as *Drosophila melanogaster, Aedes aegypti*, cell cultures and mouse models [20, 22–25]. To overcome these limitations, protein engineering yielded High-Fidelity RfxCas13d (HF-RfxCas13d), a four-mutant variant (A134V, A140V, A141V, A143V) engineered to disrupt promiscuous bystander RNA interactions. HF-RfxCas13d selectively mitigates collateral RNA cleavage and cytotoxicity. Although this is achieved at the expense of overall slightly lower on-target knockdown efficiency compared to the wild-type enzyme, HF-RfxCas13d offers an attractive, non-cytotoxic alternative for post-transcriptional gene silencing [25].

In this study we successfully established a programmable and flexible RNA depletion platform in *P. falciparum* that bypasses the need for locus-specific genomic modifications. We first engineered a stable parasite line constitutively expressing HF-RfxCas13d. We then evaluated the targeting versatility of this platform by expressing single or dual CRISPR Cas13d guide RNAs (gRNAs) of distinct length architectures (30 nt and 23 nt spacers) targeting a non-mutable gene (*atp4*) and two mutable genes (*ap2-exp* and *ap2-g*) [4]. We show that HF-RfxCas13d achieves robust and flexible, multigene transcript depletion without inducing collateral rRNA cleavage, growth defects, or developmental toxicity. Finally, we effectively applied this system to disrupt sexual conversion driven by *ap2-g*.

## 2. Methods

### Parasite culture and Transfection

*P. falciparum* asexual blood-stage parasites (*3D7 parental line* and derivative lines) were cultured in human erythrocytes at 3% haematocrit in complete RPMI 1640 medium supplemented with 25 mM HEPES, 0.5% (w/v) Albumax II, 5% (w/v) sodium bicarbonate and 0.37 mM hypoxanthine. Cultures were maintained at 37 °C under a gas mixture of 3% CO2, 1% O2, 93% N2 under static or shaking conditions (100 rpm). To minimize spontaneous sexual conversion and maintain high transfection efficiency, cultures were supplemented with 200 µM choline chloride (Sigma-Aldrich, C7527) 32 hours before to transfection. Synchronous ring-stage parasite populations were obtained by two consecutive treatments with 5% (w/v) D-sorbitol for 10 min at 37 °C spaced 12–14 h apart. Parasitaemia and developmental stages were routinely monitored by light microscopy of Giemsa-stained thin blood smears.

Transfections were performed using highly synchronous young ring-stage parasites at 5–10% parasitaemia using a Bio-Rad Gene Pulser system as previously described [26]. Drug selection was initiated 48 h post-transfection upon completion of reinvasion using 2.5 nM WR99210 (Sigma-Aldrich, SML2976). For integration transfections employing the pDC2-SpyCas9-g*Pfs47* plasmid, the drug treatment ended on day 7 and established transgenic cultures were treated with 1 µM 5-fluorocytosine (5-FC; Sigma-Aldrich, F7129) for 7 days to select against the Cas9 plasmid prior to clonal isolation. For episomal maintenance of gRNA-expressing pL6 plasmids, selection drugs were maintained continuously throughout experimental procedures. Transgenic parasite lines were cloned by limiting dilution in 96-well flat-bottom culture plates. Positive wells were expanded sequentially before cryopreservation in liquid nitrogen and genotyping PCR validation.

### Plasmid design and construction

All restriction enzymes were obtained from New England Biolabs. Plasmid DNA was propagated and maintained in *Escherichia coli* XL10-Gold Ultracompetent Cells (Agilent, 200315). The plasmids pDC2-SpyCas9-g*Pfs47* and pBS-*Pfs47*-DiCre were kindly provided by Ellen Knuepfer, pT3TS-RfxCas13d-HiFi-HA was described by Moreno-Sanchez et al [45], and pL6-CS-WR was acquired from Addgene (Addgene plasmid #122512).

To generate the pBS-*Pfs47*:Cas13d knock-in donor plasmid, pBS-*Pfs47*-DiCre was digested with SpeI and AfeI to remove one homology arm of the insertion cassette. Cohesive ends were filled using DNA Polymerase I, Klenow Fragment (New England Biolabs, M0210S). The digested plasmid backbone was gel-purified using the QIAEX II Gel Extraction Kit (Qiagen, 20021), self-ligated, and transformed into *E. coli* XL10-Gold. Successful excision was confirmed by restriction enzyme mapping. The coding sequence for HA-tagged High-Fidelity RfxCas13d (HF-RfxCas13d-HA) was PCR-amplified from pT3TS-RfxCas13d-HiFi-HA using PrimeSTAR DNA Polymerase (Takara, R045A) with primer pair P23 and P24 at an annealing temperature of 63 °C. The PCR product was purified using the MinElute PCR Purification Kit (Qiagen, 28004), digested with PstI, 5′-phosphorylated with T4 Polynucleotide Kinase (T4 PNK, New England Biolabs, M0201S), and cleaned with the MinElute kit. Concurrently, the modified pBS-*Pfs47* vector was digested with EcoRI to release genes under the *bip* promoter, bluntended with Klenow fragment, dephosphorylated using alkaline phosphatase, and digested with PstI. The vector backbone was gel-purified with QIAEX II and ligated with the digested HF-RfxCas13d-HA insert. The resulting construct was transformed into XL10-Gold *E. coli* and verified by Sanger sequencing using primers P25 and P26.

To construct the gRNA expression scaffold plasmid pL6-DR30-mutDHFR, the human dihydrofolate reductase (*hdhfr*) marker in pL6-CS-WR was first replaced with *mutDHFR* to eliminate an internal BbsI restriction site required for downstream gRNA cloning. The *mutDHFR* gene was PCR-amplified from pDC2-SpyCas9-g*Pfs47* using PrimeSTAR DNA Polymerase with primers P27 and P28, followed by MinElute purification. The pL6-CS-WR backbone and the *mutDHFR* PCR product were digested with BamHI and HindIII. The 6.1 kb digested vector fragment was gel-purified using QIAEX II, ligated with digested *mutDHFR*, and transformed into XL10-Gold *E. coli*. Correct assembly was verified by restriction enzyme digestion. Next, a synthetic double-stranded DNA fragment (GeneArt, Thermo Fisher Scientific, PRGS03KDE) containing the gRNA expression cassette with the RfxCas13d direct repeat (DR30) sequence positioned upstream of tandem BbsI cloning sites was subcloned into pL6-mutDHFR via AvrII and BstAPI restriction sites. The resulting scaffold plasmid, designated pL6-DR30-mutDHFR, was verified by Sanger sequencing using primers P29 and P30.

Individual gRNA expression plasmids were generated by inserting annealed oligonucleotide duplexes into pL6-DR30-mutDHFR. Forward and reverse single-stranded oligonucleotides corresponding to each target sequence were designed with 4-nt 5′ single-stranded overhangs (5′-AAAC-3′ on the forward strand and 5′-AAAA-3′ on the reverse strand) to complement BbsI-cleaved vector ends. Oligonucleotide pairs (1 µL each at 100 µM) were mixed with 0.5 µL T4 PNK in 1x T4 DNA Ligation Buffer, phosphorylated, and annealed in a thermocycler (37 °C for 30 min, 94 °C for 5 min, followed by cooling to 25 °C at a ramp rate of −6 °C/min). The pL6-DR30-mutDHFR vector was digested with BbsI, gel-purified using QIAEX II, and ligated with the phosphorylated oligo duplexes. Ligated products were transformed into XL10-Gold *E. coli* and minipreps were confirmed by Sanger sequencing using primer P30. Primer sequences used for vector construction and sequencing are compiled in Supplementary Table S1.

### gRNA design

Target sites for gRNA design were selected within the open reading frames of three endogenous *P. falciparum* genes: *atp4*, (PF3D7_1211900), *ap2-exp* (PF3D7_1466400) and *ap2-g* (PF3D7_1222600). Potential guide RNA target sequences were identified using the web-based design platforms *RNAtargeting* (https://rnatargeting.genomics.arcinstitute.org/) [28] for 30-nt spacer guides and *Cas13Design* (https://cas13design.nygenome.org/) [29] for 23-nt spacer guides. High-scoring target sites predicted across multiple independent algorithms were filtered to avoid potential off-targets using a BLAST search in accordance with the filters set out by Wei et al [28] in the reference RNA sequences database for *P. falciparum* isolate 3D7. For each target gene, two distinct gRNAs were selected: a 30-nt guide (designated gRNA-A) and a 23-nt guide (designated gRNA-B). Complete target sequences and guide spacer lengths configurations for all gRNAs are provided in Supplementary Table S2.

### DNA extraction and genotyping PCR

Genomic DNA was isolated using the DNeasy Blood & Tissue Kit (Qiagen, 69504). Genotyping PCRs were performed using the KAPA HiFi HotStart ReadyMix PCR Kit (Roche, 09420398001). Reactions contained 100 ng of genomic DNA template, 0.5 µM of each forward and reverse primer, 1 mM supplementary MgCl2, and 1x KAPA HiFi HotStart ReadyMix (25 µL). Thermocycling conditions comprised an initial denaturation at 95 °C for 3 min; 30 cycles of 98 °C for 20 s, target-specific annealing temperature for 15 s (Supplementary Table S1), and 72 °C for 1 min 35 s; followed by a final extension at 72 °C for 1 min 35 s. PCR products were resolved by 1.5% agarose gel electrophoresis and visualized under ultraviolet light.

### Protein extraction, quantification and western blotting

Saponin-isolated whole parasite pellets were prepared for protein expression analysis. Cultured parasites (5 mL) were washed twice with 25 mL cold PBS (centrifuged at 800 g for 5 min). Erythrocytes were lysed by resuspending in 0.05% (w/v) saponin in PBS and rocking gently for 3 min at room temperature until turbidity cleared. Released parasites were pelleted by centrifugation at 1800 g for 10 min, washed with PBS, and resuspended in 1 mL cold PBS containing 1x EDTA-free Protease Inhibitor Cocktail (PIC; Roche, 05892791001). Parasites were pelleted at 16,000 g for 1 min. Parasite pellets were solubilized in Parasite Lysis Buffer (4% SDS, 0.5% Triton X-100 in 1x PBS) containing 1x PIC. Lysates were centrifuged at 16,000 g for 5 min at 4 °C to remove insoluble debris, and supernatants were transferred to clean tubes.

Total protein concentration was quantified using the DC Protein Assay Kit (Bio-Rad, 5000112) according to the manufacturer’s detergent-compatible protocol. Bovine serum albumin (BSA) standards (0.156–10.0 mg/mL) were prepared in Parasite Lysis Buffer. Samples, standards, and buffer blanks (5 µL per well in duplicate) were loaded into a 96-well plate. Absorbances were measured at 750 nm on a Molecular Devices VERSAmax Tunable Microplate Reader running SoftMax Pro. Sample protein concentrations were calculated from the BSA linear standard curve. Quantified extracts were diluted with 4x Laemmli Sample Buffer (Bio-Rad) with 10% (v/v) β-mercaptoethanol to a 1x final concentration and denatured at 95 °C for 5 min prior to electrophoresis or storage at −20°C.

Equivalent protein mass (33 μg) was separated on TGX™ FastCast™ acrylamide gels (Bio-Rad, 1610173) at 200 V for 40 min. Precision Plus Protein All Blue Standards (Bio-Rad, 1610373; 8 µL) served as the molecular weight marker. Proteins were transferred onto methanol-pre-wetted PVDF membranes (Bio-Rad, 1620174) at 325 mA for 1–2 h at 4 °C. Membranes were blocked with 5% (w/v) skim milk in TBST (1x TBS with 0.1% Tween 20) for 30–60 min at room temperature with agitation. Primary antibody incubations were performed overnight at 4 °C with agitation in blocking buffer using rat anti-HA (1:1,000; Roche, Cat. No. 11867423001; target protein ∼113 kDa) and rabbit anti-*Plasmodium* aldolase (1:1,000; Abcam, ab207494; loading control, 40 kDa). Membranes were washed three times with TBST and incubated for 1 h at room temperature in the dark with fluorophore-conjugated secondary antibodies diluted 1:10,000 in blocking buffer: Goat anti-Rat IgG (H+L) Cross-Adsorbed Alexa Fluor 680 (Thermo Fisher Scientific, A21096; red channel 700 nm) and Goat anti-Rabbit IgG H&L IRDye 800CW pre-absorbed (Abcam, ab216773; green channel 800 nm). Following three washes in TBST and a final wash in TBS protected from light, fluorescent signal was captured using an Odyssey CLx Near-Infrared Imaging System (LI-COR Biosciences) at 700 nm and 800 nm wavelengths.

### Quantitative western blot and densitometric analysis

Digital image processing and quantitative densitometric analysis of HF-RfxCas13d expression were performed using Fiji/ImageJ (NIH). Non-saturated 32-bit TIFF images acquired on the Odyssey CLx scanner were converted to 8-bit grayscale matrices. Images were inverted so that protein bands represented positive high-intensity pixel values above a dark background. Local background subtraction was conducted using the Gels analysis module. Equal rectangular regions of interest encompassing the migration front of each protein band across developmental stages (ring, trophozoite, and schizont) were delineated, translated across lanes, and plotted as 1D intensity profile plots (Plot Lanes).

Baseline local background noise was eliminated by drawing a straight line across the base of each peak. The background-subtracted integrated density of each band was calculated using the Wand Tool. Because all wells were loaded with equal total parasite protein mass as determined by the DC protein assay, the background-subtracted integrated density values provided direct measurements of protein abundance. Signal intensities were not normalized to an internal housekeeping protein because standard loading markers, including *PfAldolase*, exhibit stage-dependent expression variability across the developmental cycle (Supplementary Fig. S2d, e) Stage-specific expression changes across the IDC were expressed as Fold Change relative to ring-stage baseline values.

Statistical significance of stage-specific HF-RfxCas13d protein variations across independent biological replicates (n = 3) was evaluated using two-tailed Student’s t-tests in GraphPad Prism 10.

### Immunofluorescence microscopy

Parasite-infected erythrocytes at >5% parasitaemia were collected and seeded onto round glass coverslips placed in 24-well culture plates. Cells were fixed with 4% (v/v) formaldehyde, 0.0075% (v/v) glutaraldehyde in PBS for 20 min at 37 °C. Free aldehyde groups were quenched by incubating with 30 mM glycine. Parasites were permeabilized with 0.1% (v/v) Triton X-100 for 20 min and blocked with 3% (w/v) BSA for 45 min at room temperature. HF-RfxCas13d-HA was detected using rat anti-HA (1:600; Roche, Cat. No. 11867423001) for 1 h at room temperature and with secondary fluorophore-conjugated antibody (Goat anti-Rat IgG Cross-Adsorbed Alexa Fluor 680, 1:1,000; Invitrogen, A-21096) for 1 h at room temperature). Parasite nuclei were stained with 4’,6-diamidino-2-phenylindole (DAPI, 2 μg/mL final) for 5 min at room temperature. Coverslips were mounted onto glass slides using Vectashield mounting medium (Vector Laboratories, H-1700). Fluorescence imaging was performed on a Leica DMi8 inverted microscope with 100X objective, and raw images were analysed using Fiji/ImageJ (NIH).

### Growth analysis

Asexual blood-stage growth rates were evaluated in tightly synchronized parasite cultures. Ring-stage parasites were seeded at an initial parasitaemia of 0.1% and 2% haematocrit in 6-well plates in biological triplicates for each condition. Parasitaemia was quantified by flow cytometry at 24 h (Cycle 0), 72 h (Cycle 1), and 96 h (Cycle 2). At each time point, a sample was fixed in 0.2% (v/v) glutaraldehyde for 1 h at room temperature. Fixed samples were washed and stored in fresh PBS at 4 °C. Prior to acquisition, fixed cells were stained with 8 µM Hoechst 33342 (Invitrogen, H1399) for 10 min at room temperature. An unstained parasite sample was processed in parallel as a background control. Parasites were acquired on a BD FACSymphony flow cytometer. The percentage of infected red blood cells was determined by gating on Hoechst-positive populations across 50,000 recorded events per sample using FlowJo (version 10.8.1).

Statistical significance of parasite growth variations across independent biological replicates (n = 3) was evaluated using two-way ANOVA with multiple comparisons in GraphPad Prism 10.

### RNA extraction, cDNA synthesis and qPCR

Total RNA was isolated from highly synchronized blood-stage parasite cultures at specific developmental stages. Infected erythrocytes were pelleted, lysed by adding 10 volumes of TRIzol reagent (Invitrogen) and incubated for 5 min at 37 °C before storage at −80 °C. Samples were thawed on ice, mixed with 0.2 volumes of chloroform per volume of TRIzol and incubated on ice for 5 min. Phase separation was achieved by centrifugation at 4,500*xg* for 15 min at 4 °C brake off. The upper aqueous phase (80% volume) was transferred to an RNase-free tube, mixed with 1 volume of 100% ethanol, and total RNA was extracted using Direct-zol RNA Microprep column (Zymo Research, R2060) with incolumn DNase I digestion. RNA concentration and purity were measured using a NanoDrop spectrophotometer. RNA integrity was verified by resolving 100 ng of RNA mixed with Ember500 RNA Prestain Loading Dye (Biotium, 41032) on a 1% agarose gel.

For cDNA synthesis, total RNA was reverse-transcribed using the RevertAid First Strand cDNA Synthesis Kit (Thermo Scientific, K1622) with random hexamer primers according to the manufacturer’s instructions. Quantitative real-time PCR (RT-qPCR) was performed using the KAPA SYBR FAST qPCR Kit (Roche, KK4601) on a Bio-Rad CFX96 Real-Time PCR Detection System with technical triplicates. Reaction conditions consisted of initial denaturation at 95 °C for 3 min, followed by 40 cycles of 95 °C for 3 s, 54°C for 20 s, and extension at 60 °C for 30 s. 54°C annealing temperature was applied to all RT-qPCR primer pairs after melt-curve and primer efficiency validation for single amplicon quantification. Target gene transcript levels were normalized against a normalization factor derived from the arithmetic mean Cq values of two internal reference genes: *serine-tRNA ligase* (*serr*, PF3D7_0717700) and *ubiquitin-conjugating enzyme E2* (*uce*, PF3D7_0812600). Relative fold changes in target mRNA abundance were calculated using the ΔΔCq method.

Statistical significance of transcript variations across independent biological replicates (n = 3) was evaluated using two-tailed Student’s t-tests in GraphPad Prism 10.

### Sexual conversion induction and assays

Gametocytogenesis was induced using choline depletion as described [30]. Briefly, parasites were initially grown under shaking conditions (100 rpm) at 37 °C in complete medium containing 2 mM choline chloride for at least two cycles and parasitaemia maintained below 2%. For the induction phase (Cycle 0), ring-stage cultures were sorbitol-synchronized, adjusted to 1.0–2.5% parasitaemia at 3% haematocrit, and washed with medium to remove residual choline. The culture was split into two parallel conditions in 6-well plates under static conditions: Gametocyte-repressed control (+ choline) and Gametocyte-induced condition (-choline), cultured in complete medium with or without 2mM choline chloride, respectively.

In the subsequent cycle (Cycle 1), when parasites reached ring stage, total parasitaemia was determined from Giemsa-stained thin smears, and complete medium with 50 mM *N*-acetyl-D-glucosamine (GlcNAc; Sigma-Aldrich, A3286) and 2 mM choline was added to selectively eliminate asexual blood-stage parasites. Media containing 50 mM GlcNAc were renewed daily for 3–4 days. On Day 4, thin blood smears were prepared to count mature gametocytes. The Sexual Conversion Rate (SCR) was calculated by dividing the gametocytemia on Day 4 over the initial parasitaemia of the culture at the ring stage.

Statistical significance of SCR variations across independent biological replicates (n = 3-4) was evaluated using one-way ANOVA with multiple comparisons in GraphPad Prism 10.

## 3. Results

### Generation of a stable marker-free HF-RfxCas13d expressing parasite line

To establish a programmable RNA depletion platform in *P. falciparum*, we engineered a parasite line constitutively expressing High-Fidelity RfxCas13d (HF-RfxCas13d) (Fig. 1a). The expression cassette encoding C-terminally HA-tagged HF-RfxCas13d under the control of the *bip* promoter (5′ UTR) and *cam* 3′ UTR was integrated into the dispensable *pfs47* locus of the 3D7 parental line via CRISPR-Cas9-mediated homologous recombination, following a marker-free approach adapted from Knuepfer et al.[31].

**Figure 1.**
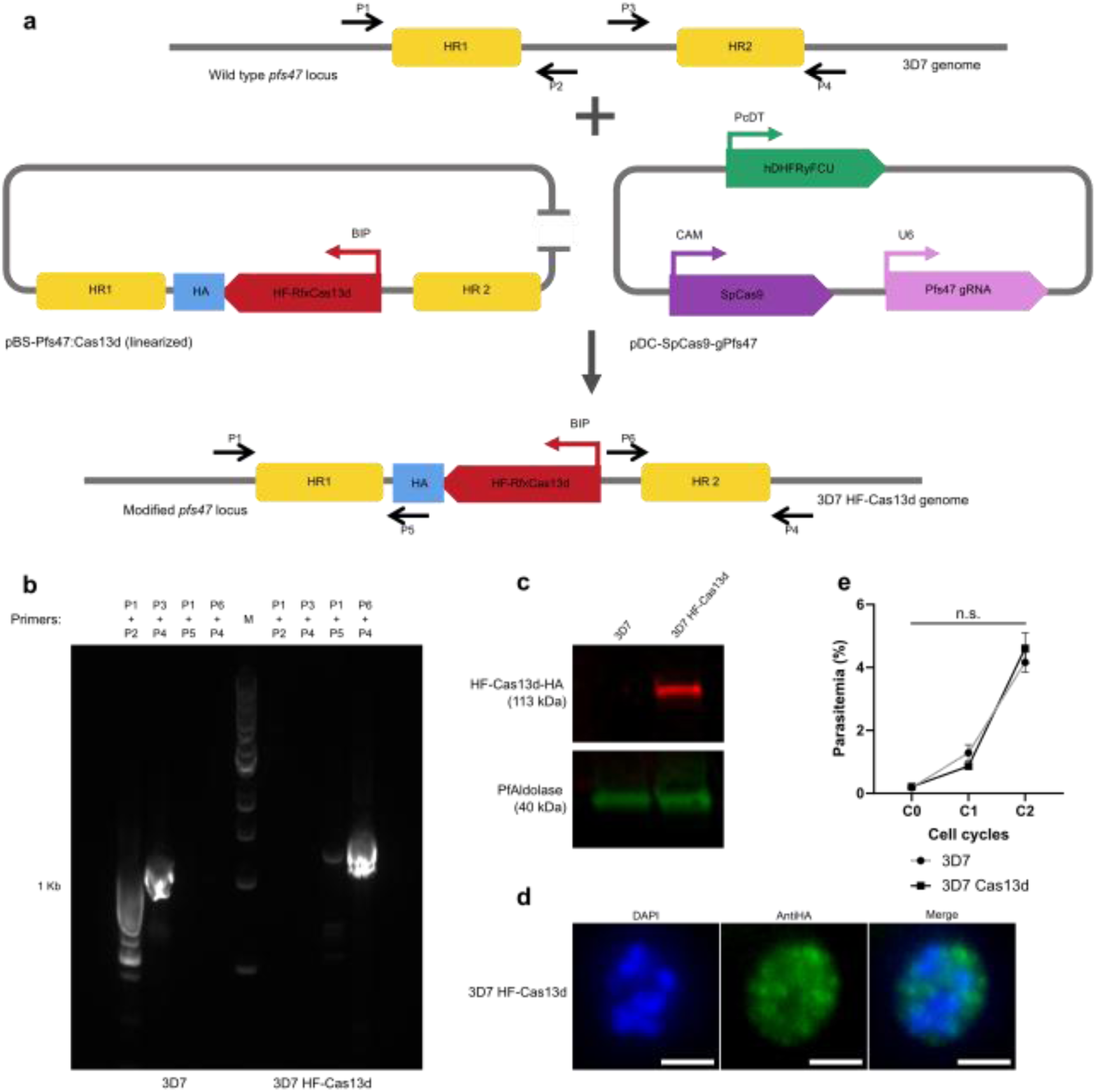
Generation and characterization of 3D7 HF-Cas13d parasite line. (**a**) Schematic representation of the wild-type *pfs47* locus, the CRISPR-Cas9 plasmids used for knock-in of the HF-RfxCas13d expression cassette, and the modified HF-RfxCas13d-expressing *pfs47* locus. Black arrows denote primer binding sites used for genotyping. (**b**) Genotyping PCR validation of HF-RfxCas13d expression cassette integration in the 3D7 HF-Cas13d clone and the 3D7 parental line. Primer pairs P1+P2 and P3+P4 amplify the wild-type *pfs47* locus, whereas primer pairs P1+P5 and P6+P4 amplify specifically the modified locus. M: molecular weight marker. (**c**) Western blot analysis detecting HF-RfxCas13d expression (anti-HA) in the 3D7 parental line and 3D7 HF-Cas13d. PfAldolase serves as a loading control. Cropped blots are displayed; full-length blots are provided in Supplementary Fig. S2. (**d**) Immunofluorescence assay of 3D7 HF-Cas13d using DAPI (blue) and anti-HA antibody (green) demonstrating HF-RfxCas13d expression. Scale bar, 2 µm. (**e**) Parasite growth of the 3D7 parental line and 3D7 HF-Cas13d clone after a 96 h growth assay. No statistically significant difference was observed (p=0.8990, two-way ANOVA with multiple comparisons, n=3). Error bars represent Mean ± SEM.

We genotyped eight isolated clonal lines and confirmed targeted integration of the HF-RfxCas13d cassette at the *pfs47* locus in all evaluated clones. Diagnostic amplicons evidenced specific amplification of 5′ and 3′ integration junction fragments, alongside the concurrent loss of the wild-type fragments (Fig. 1b and Supplementary Fig. S1). We selected Clone D4 for subsequent functional characterization and was designated 3D7 HF-Cas13d.

Expression of the HF-RfxCas13d protein in the 3D7 HF-Cas13d clonal line was verified by western blot analysis using an anti-HA antibody, which detected a single band corresponding to the expected molecular weight (∼113 kDa) that was absent in the 3D7 parental line (Fig. 1c and Supplementary Fig. S2a). Immunofluorescence microscopy confirmed HF-RfxCas13d protein expression and demonstrated its cytosolic distribution in 3D7 HF-Cas13d parasites (Fig. 1d). To assess whether constitutive expression of HF- RfxCas13d induces cellular toxicity during blood-stage development, we evaluated the asexual replication kinetics over 96 h (two intraerythrocytic developmental cycles). The parasite growth of 3D7 HF-Cas13d showed no statistically significant difference compared to the 3D7 parental line (adjusted p = 0.8990; Fig. 1e), confirming that constitutive HF-RfxCas13d expression does not impair asexual parasite proliferation.

### HF-RfxCas13d system allows RNA depletion of endogenous genes

To evaluate the versatility and efficacy of the 3D7 HF-Cas13d line as a programmable RNA depletion platform in *P. falciparum*, we targeted four endogenous parasite genes: a non-mutable: *atp4* (PF3D7_1211900); and two mutable: *ap2-exp* (PF3D7_1466400) and *ap2-g* (PF3D7_1222600), as previously stated by High-throughput piggyBac transposon insertional mutagenesis [4]. Given the lack of data on *P. falciparum*, we used described algorithms for eukaryotic RfxCas13d RNA design. For each target gene, two distinct CRISPR Cas13d guide RNAs (gRNAs) were designed: a 30-nt guide spacer (gRNA-A) using RNAtargeting [28] and a 23-nt guide spacer (gRNA-B) using Cas13Design (Fig. 2a-b and Table S2) [29]. All gRNA cassettes were individually cloned into separate customized scaffold plasmid pL6-DR30-mutDHFR under the control of the PfU6 promoter. To assess simultaneous targeting, gRNAs for *atp4* (gATP4 A and gATP4 B) or *ap2-exp* (gAP2-EXP A and gAP2-EXP B) were co-transfected into 3D7 HF-Cas13d cultures.

**Figure 2.**
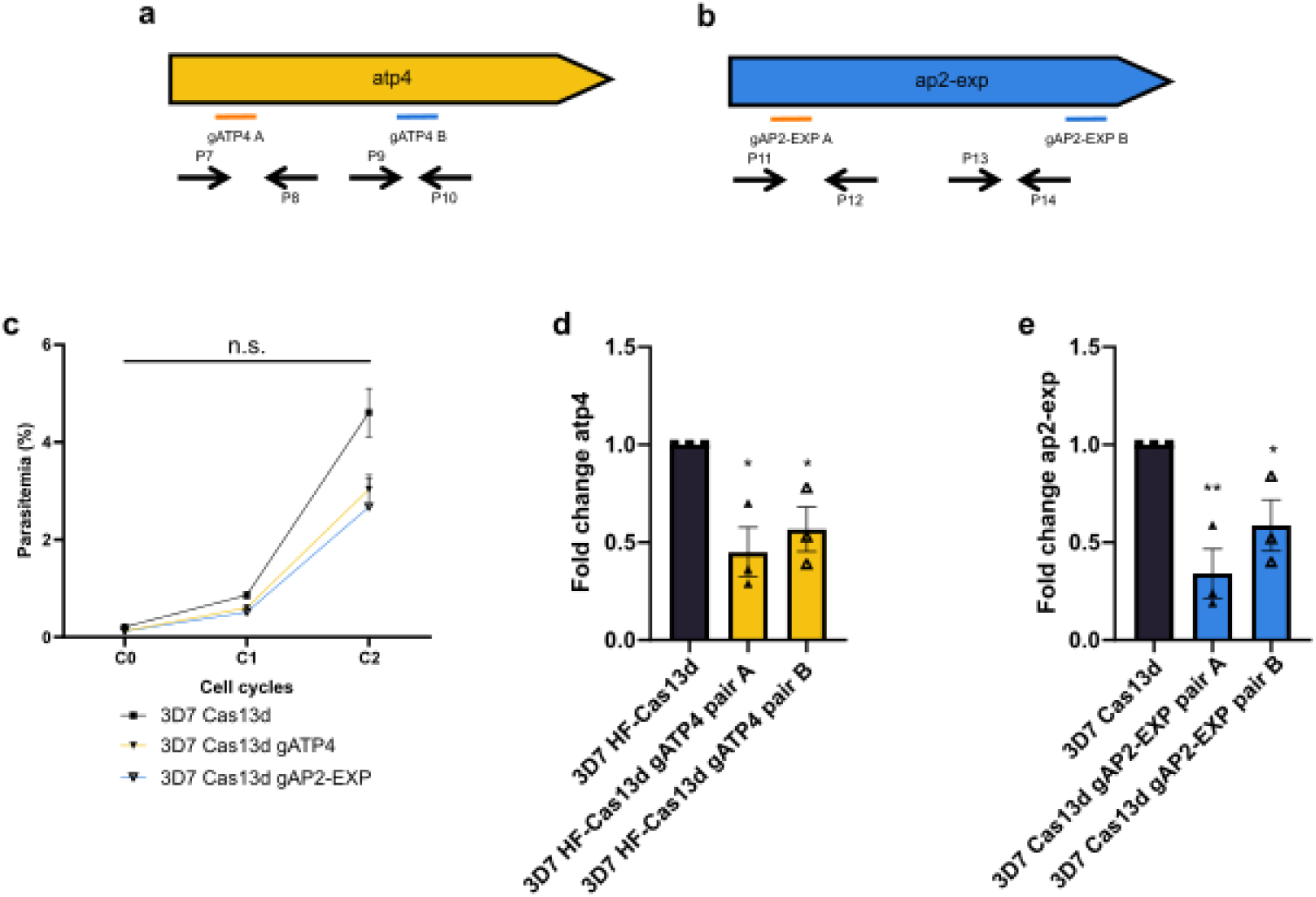
HF-RfxCas13d-mediated target RNA depletion of endogenous parasite genes. (**a–b**) Schematic representations of *atp4* (**a**) and ap2-exp (**b**) open reading frames. Lines indicate the relative target positions of gRNA-A (30 nt, orange) and gRNA-B (23 nt, blue). Black arrows represent RT-qPCR primer pairs designed adjacent to or flanking the target site. Plasmids expressing gRNAs for *atp4* or *ap2-exp* were cotransfected as dual-gRNA combinations in single 3D7 HF-Cas13d cultures. (**c**) Parasite growth after 96 h comparing 3D7 HF-Cas13d with targeted cultures: 3D7 HF-Cas13d gATP4 (adjusted p=0.2875, n.s.), and 3D7 HF-Cas13d gAP2-EXP (adjusted p=0.1661, n.s.). No statistically significant differences were observed (two-way ANOVA with multiple comparisons, n=3). (**d–e**) Target mRNA expression analysis by RT-qPCR. Fold Change values were determined using the ΔΔCq method normalized to *serr* and *uce* reference mRNAs. Significant target RNA depletion was detected across all targets compared to the 3D7 HF-Cas13d control: 3D7 HF-Cas13d gATP4 with primer pair A (p=0.0122) and pair B (p=0.0191), and 3D7 HF-Cas13d gAP2-EXP with primer pair A (p=0.0063) and pair B (p=0.0346). *p<0.05, **p<0.01, two-tailed t-test (n=3). Error bars represent Mean ± SEM.

Next, we investigated whether activation of HF-RfxCas13d by target-bound gRNAs induced non-specific bystander RNA cleavage (collateral activity) or general cellular toxicity in *P. falciparum*. Analysis of asexual replication kinetics over 96 h revealed no statistically significant differences in parasite growth between the unguided 3D7 HF-Cas13d control line and any of the gRNA-expressing parasite lines (p > 0.05 across all comparisons; Fig. 2c). Specifically, parasitaemia were not significantly altered in 3D7 HF-Cas13d gATP4 (adjusted p = 0.2875), and 3D7 HF-Cas13d gAP2-EXP (adjusted p = 0.1661). Furthermore, total RNA gel electrophoresis confirmed the integrity of 28S and 18S ribosomal RNA bands across all conditions, suggesting the absence of non-specific bystander transcript processing or collateral rRNA degradation (Supplementary Fig. S3a). Altogether, these results strongly suggest that HF-Cas13d expression in *P. falciparum* did not cause toxic effects.

To determine CRISPR-HF-Cas13d activity in *P. falciparum*, we measured relative transcript abundance represented by fold change (FC) by RT-qPCR using amplicons situated adjacent to or spanning the gRNA binding sites. Dual gRNA delivery targeting *atp4* yielded substantial transcript depletion measured by two independent diagnostic primer pairs, showing a 55% reduction with primer pair A (p = 0.0122) and a 43% reduction with primer pair B (p = 0.0191) (Fig. 2d). Similarly, co-expression of two gRNAs targeting *ap2-exp* produced significant mRNA knockdown, resulting in a 66% reduction measured by primer pair A (p = 0.0063) and a 41% reduction measured by primer pair B (p = 0.0346) (Fig. 2e). Together, these results show that the HF-RfxCas13d platform enables targeted and significant RNA depletion of multiple endogenous *P. falciparum* genes (ranging from 41% to 66%) without inducing collateral rRNA cleavage or off-target growth inhibition.

### RNA depletion of *ap2-g* by HF-RfxCas13d impairs sexual conversion

To evaluate whether the High-Fidelity RfxCas13d transcript depletion can functionally disrupt well-known phenotypes, we targeted *ap2-g* (PF3D7_1222600), the master transcriptional regulator of gametocytogenesis in *P. falciparum* [32, 33]. To evaluate different guide delivery strategies, two single gRNA expression plasmids targeting distinct sites within the *ap2-g* open reading frame -gAP2-G A (30 nt) and gAP2-G B (23 nt)-were independently transfected into 3D7 HF-Cas13d parasites and selected with WR99210 (Fig. 3a and Supplementary Table S2). Evaluation of asexual replication kinetics over 96 h revealed no statistically significant differences in parasitaemia between the unguided 3D7 HF-Cas13d parent line and either 3D7 HF-Cas13d gAP2-G A (p = 0.9939) or 3D7 HF-Cas13d gAP2-G B (p = 0.8893) (Fig. 3b). Total RNA gel electrophoresis further verified intact 28S and 18S ribosomal RNA bands across both ring and schizont stages, implying no substantial non-specific bystander rRNA cleavage upon *ap2-g* targeting (Supplementary Fig. S3b).

**Figure 3.**
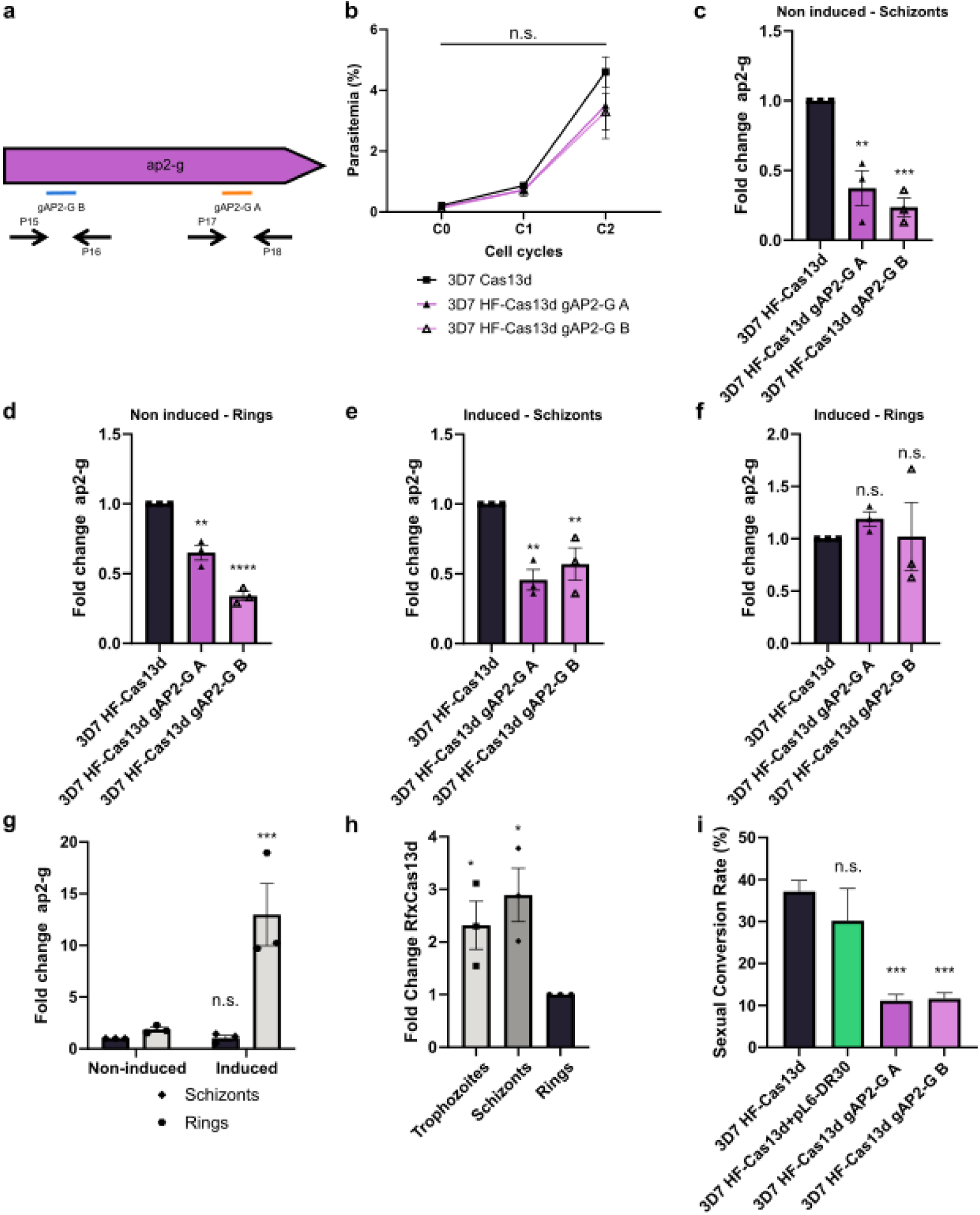
*ap2-g* RNA depletion by HF-RfxCas13d impairs sexual conversion rate in *P. falciparum*. (a) Schematic representation of the *ap2-g* open reading frame depicting target positions of gAP2-G A (30 nt, orange) and gAP2-G B (23 nt, blue), alongside RT-qPCR primer pairs (black arrows). Plasmids expressing gAP2-G A and gAP2-G B were transfected into independent 3D7 HF-Cas13d cultures. (b) Parasitaemia after 96 h of 3D7 HF-Cas13d and *ap2-g*-targeted cultures. No significant differences were observed (gAP2-G A: adjusted p=0.9939; gAP2-G B: adjusted p=0.8893; two-way ANOVA with multiple comparisons, n=3). (c–f) RT-qPCR analysis of *ap2-g* mRNA expression in non-induced schizonts (c), non-induced rings (d), choline-induced schizonts (e), and cholineinduced rings (f). Fold Change values were calculated using the ΔΔCq method normalized to *serr* and *uce* relative to 3D7 HF-Cas13d. Non-induced schizonts: gAP2-G A (p=0.0076), gAP2-G B (p=0.0003). Non-induced rings: gAP2-G A (p=0.0027), gAP2-G B (p<0.0001). Induced schizonts: gAP2-G A (p=0.0017), gAP2-G B (p=0.0207). Induced rings: gAP2-G A (p=0.0518), gAP2-G B (p=0.9542). Two-tailed t-test (n=3). (g) Analysis of *ap2-g* mRNA relative levels in 3D7 HF-Cas13d across developmental stages and induction condition using primer pair A, normalized to non-induced schizonts. Two-way ANOVA revealed significant differences between induced and non-induced rings (p=0.0008), but not schizonts (p=0.9867). (h) HF-RfxCas13d protein quantification across the IDC. Fold Change values determined by western blot (anti-HA) relative to rings demonstrate increased expression in trophozoites (p=0.0440) and schizonts (p=0.0204; two-tailed t-test, n=3). (i) Sexual Conversion Rates (%) following choline depletion in 3D7 HF-Cas13d (n=4), control 3D7 HF-Cas13d+pL6-DR30 (n=3), and *ap2-g*-targeted cultures A and B (n=4). Significant drops were detected in 3D7 HF-Cas13d gAP2-G A (adjusted p=0.0005) and 3D7 HF-Cas13d gAP2-G B (adjusted p=0.0006) relative to 3D7 HF-Cas13d, whereas control 3D7 HF-Cas13d+pL6-DR30 showed no significant change (adjusted p=0.4195; one-way ANOVA with multiple comparisons). *p<0.05, **p<0.01, ***p<0.001, ****p<0.0001. Error bars represent Mean ± SEM.

Because *ap2-g* transcription is markedly upregulated during sexual commitment [7], we quantified relative *ap2-g* transcript abundance across distinct developmental stages (schizonts and post-commitment rings) under both gametocyte-repressed (+ choline) and gametocyte-induced (-choline) conditions using RT-qPCR (Supplementary Fig. S4). We induced gametocytogenesis by choline depletion following the protocol described by Portugaliza et al. [30]. In non-induced parasite populations, both gRNAs mediated significant target mRNA depletion relative to 3D7 HF-Cas13d. In non-induced schizonts, *ap2-g* mRNA was depleted by 63% with gAP2-G A (p = 0.0076) and by 76% with gAP2-G B (p = 0.0003) (Fig. 3c); in non-induced rings, *ap2-g* transcript abundance decreased by 35% with gAP2-G A (p = 0.0027) and by 66% with gAP2-G B (p < 0.0001) (Fig. 3d). Under choline-depleted, sexual-inducing conditions, significant mRNA depletion was likewise observed in schizonts, showing a 54% reduction with gAP2-G A (p = 0.0017) and a 43% reduction with gAP2-G B (p = 0.0207) (Fig. 3e). On the other hand, in post-commitment ring-stage parasites, no statistically significant differences in *ap2-g* transcript abundance were detected for either gAP2-G A (p = 0.0518) or gAP2-G B (p = 0.9542) compared to the 3D7 HF-Cas13d control (Fig. 3f). Altogether, these results indicate that HF-RfxCas13d mediates robust target RNA depletion of *ap2-g* across multiple developmental stages but fails to reduce transcript abundance during the post-commitment ring stage when target mRNA expression peaks.

To investigate the transcriptomic and protein expression dynamics underlying this stage-specific knockdown pattern, we quantified baseline *ap2-g* mRNA expression in 3D7 HF-Cas13d parasites normalized to non-committed schizonts-stage levels, alongside HF-RfxCas13d protein abundance across the Intraerythrocytic Developmental Cycle (IDC) (Fig. 3g, h). Two-way ANOVA revealed a sharp increase in *ap2-g* transcript abundance in induced rings relative to non-induced rings (p = 0.0008), whereas no significant difference in baseline transcript levels was observed between induced and non-induced schizonts (p = 0.9867) (Fig. 3g). Simultaneously, western blot analysis of HA-tagged HF-RfxCas13d across synchronized parasite stages showed that HF-RfxCas13d protein expression is stage-dependent, displaying significant fold-increases in trophozoites (p = 0.0440) and schizonts (p = 0.0204) relative to ring-stage baseline levels (Fig. 3h and Supplementary Fig. S2b, c).

To measure the phenotypic and functional consequences of the knockdown *ap2-g* mutants, we quantified the Sexual Conversion Rate (SCR) following choline starvation (Fig. 3i) [5]. Target depletion of *ap2-g* caused a robust and statistically significant decrease in gametocyte formation relative to the unguided parent line (3D7 HF-Cas13d SCR = 37.15±2.65%), with 3D7 HF-Cas13d gAP2-G A and 3D7 HF-Cas13d gAP2-G B achieving SCRs of 11.14±1.46% (adjusted p=0.0005) and 11.62±1.39% (adjusted p=0.0006), respectively, as determined by one-way ANOVA with multiple comparisons. In contrast, the control line targeting an unrelated gene exhibited an SCR of 30.18±7.68%, showing no statistically significant difference from 3D7 HF-Cas13d (adjusted p=0.4195) (Fig. 3i). These results indicate that HF-RfxCas13d-mediated specific depletion of *ap2-g* transcripts effectively reduces sexual conversion in *P. falciparum*.

## 4. Discussion

In this study, we successfully established a stable *P. falciparum* parasite line constitutively expressing High-Fidelity RfxCas13d (3D7 HF-Cas13d). By delivering episomal guide cassettes targeting three endogenous genes with distinct mutability profiles [4] we achieved targeted transcript depletion ranging from 41% to 76%. This RNA depletion occurred without detectable collateral effects and preserving intraerythrocytic asexual replication across all tested conditions. Applying HF-RfxCas13d to the master transcriptional regulator of gametocytogenesis (*ap2-g*) functionally disrupted sexual development from a Sexual Conversion Rate of 37.15 % in the unguided parent line to ∼11% in *ap2-g*-targeted cultures.

The advantages and limitations of our 3D7 HF-Cas13d tool are best evaluated when benchmarked against the conditional toolbox available for *P. falciparum*. At the DNA level, DiCre-mediated recombination achieves >90% excision efficiency within a single cycle [14, 15, 31, 35], but deletion of essential loci can result in parasite lethality, precluding long-term phenotypic analyses. At the protein level, destabilizing domains (DDs), auxin-inducible degrons (AID), and knock-sideways (KS) enable post- translational regulation; however, they require continuous small-molecule stabilization or induction, may exhibit basal leakiness, can disrupt protein topology, and are unsuitable for secreted or membrane-anchored proteins lacking access to cytosolic degradation machinery [9, 12, 36, 37]. At the RNA level, the glmS ribozyme typically produces 50- 90% transcript and protein depletion following glucosamine treatment, although this may be insufficient to generate phenotypes for low-turnover or highly abundant essential proteins [5, 11]. The TetR-DOZI system can achieve stronger repression, reaching up to 100-fold knockdown when aptamers are placed in both the 5′ and 3′ UTRs [38], but requires complex multi-step locus engineering that remains vulnerable to recombination and truncation events despite recent improvements [10, 36, 37]. Importantly, all current conditional systems require target-specific genomic modification to introduce tags, degrons, ribozymes, aptamers, or artificial introns. In contrast, HF-RfxCas13d bypasses direct modification of the target locus entirely. Although its maximal depletion (41-76%) is lower than that achieved with dual-aptamer TetR-DOZI, it provides a rapid and flexible knockdown strategy.

Targeting *ap2-g* showed that knockdown efficiency depends on the combined effects of stage-specific Cas13d abundance and endogenous target transcript dynamics during intraerythrocytic development. Lower RfxCas13d abundance in ring stages, consistent with the globally repressed transcriptional state of early intraerythrocytic development [40–42], likely limits knockdown efficacy at the time when sexual commitment induces a sharp increase in ap2-g transcript abundance through a positive autoregulatory feedback loop [32–34, 43]. Under these conditions, the surge in target mRNA may exceed the available HF-RfxCas13d pool, explaining the absence of measurable transcript depletion in committed rings. Nevertheless, partial ap2-g depletion during the preceding schizont stage was sufficient to reduce sexual conversion by ∼70%, underscoring the high sensitivity of commitment to AP2-G dosage. This phenotype is consistent with previous studies showing that genetic disruption or conditional depletion of PfAP2-G abolishes gametocyte formation [32].

In contrast to ap2-g, HF-RfxCas13d-mediated depletion of the essential blood-stage gene atp4 did not produce detectable growth defects, differing from previous TetR-DOZI studies [10]. This likely reflects the fact that phenotypic outcomes depend on the abundance and turnover of the target protein [5, 8, 44]. For highly abundant or stable proteins, the level of depletion achieved (31-55%) may be insufficient to impair parasite growth. Nevertheless, such partial knockdown could be advantageous for generating hypomorphic states that enable the study of essential genes without the lethality associated with complete gene disruption [19, 39]. As no target-specific phenotypic assays were performed, we cannot exclude the presence of sublethal physiological effects. Beyond target-specific requirements, neither spacer length nor guide multiplexing appeared to substantially influence knockdown efficiency. To explore guide design parameters in *P. falciparum*, we tested single 30 nt spacers, single 23 nt spacers, and dual-guide delivery, as multiplexed targeting has been reported to enhance Cas13- mediated cleavage [17, 19]. Although dual-guide delivery achieved higher depletion of ap2-exp, single 23 nt and 30 nt guides produced comparable knockdown of ap2-g, consistent with recent Cas13 studies [27]. Together, these results suggest that HF- RfxCas13d can accommodate diverse guide architectures without an obligatory requirement for multiplexing. The lower depletion achieved in this study may reflect the catalytic trade-off of the High-Fidelity RfxCas13d variant. While the mutations introduced to suppress collateral cleavage can reduce on-target activity [25, 27, 45], the 3D7 HF- Cas13d platform maintained target-dependent phenotypic effects without evidence of non-specific toxicity or growth impairment. Nevertheless, the absence of a control for collateral RNA cleavage prevents definitive exclusion of off-target endoribonuclease activity. Future applications requiring stronger knockdown could evaluate wild-type RfxCas13d, particularly when targeting low-abundance transcripts for which collateral activation is absent or biologically negligible [23, 27]. Beyond individual gene characterization, the 3D7 HF-Cas13d line could offer broader utility for high-throughput target screening and multiplexed gene knockdowns via the co-transfection of multiple gRNA expression plasmids [17, 27–29, 45]. Unlike CRISPR-Cas9 platforms, Cas13d does not require the design and co-delivery of custom homology repair templates for every targeted locus. Furthermore, the current constitutive expression of HF-RfxCas13d can potentially enable functional genomics across the complete *P. falciparum* life cycle, providing a stable background to interrogate gene function during sexual differentiation, mosquito-stages and liver-stage development [31, 35]. However, this setup is unsuitable for maintaining lines with target knockdowns that cause acute asexual lethality. This limitation could be worked around by incorporating inducible control to Cas13d expression, but further validations would be needed. Additionally, *pfs47*-disrupted parasites cannot successfully infect *Anopheles gambiae* mosquitoes [46–48]. To overcome this vector transmission limitation in future studies involving *A. gambiae*, the HF-RfxCas13d expression cassette could be integrated into alternative, dispensable genomic loci that do not impair mosquito infectivity, such as *p230p* [31, 46].

In conclusion, the 3D7 HF-Cas13d platform provides a proof-of-principle of a scalable, flexible, and non-toxic addition to the *P. falciparum* genetic toolbox. By enabling targeted, flexible and specific transcript depletion without requiring genomic modification of endogenous loci or inducing collateral RNA toxicity, this platform advances functional genomics and therapeutic target validation in malaria parasites.

## 5. Author statements

### 5.1. Author contributions

- **Rafael Serrano-Durán** — Conceptualization, Methodology, Validation, Formal analysis, Investigation, Data curation, Writing — original draft, Visualization.

- **Diana López-Farfán** — Conceptualization, Methodology, Formal analysis, Resources, Writing — review & editing, Supervision.

- **Miguel A. Moreno-Mateos** — Conceptualization, Methodology, Resources, Writing

— review & editing.

- **Elena Gómez-Díaz** — Conceptualization, Methodology, Formal analysis, Writing — review & editing, Supervision, Funding acquisition.

### 5.2. Conflicts of interest

The authors declare that there are no conflicts of interest

### 5.3. Funding information

This work was supported by the Spanish Ministry of Science and Innovation (grant no. PID2019-111109RB-I00), and by La Caixa Foundation—Health Research Program (grant no. HR20-00635). R.S-D. was funded by FPU fellowship (grant no. FPU20/04280) of the Spanish Ministry of Universities. The Moreno-Mateos lab is supported by grant PID2024-158755NB-I00 funded by MICIU/AEI/10.13039/501100011033 and FEDER, UE and co-financed by the Spanish Ministry of Science and Innovation with funds from the European Union NextGenerationEU (PRTR-C17.I1) and the Regional Ministry of University, Research and Innovation of the Autonomous Community of Andalusia within the framework of the Biotechnology Plan applied to Health. The CABD is an institution funded by Pablo de Olavide University, Consejo Superior de Investigaciones Científicas (CSIC), and Junta de Andalucía.

## Acknowledgements

The authors gratefully acknowledge Dr Ellen Knuepfer (The Royal Veterinary College, UK) for providing plasmids and technical advice, Dr. Manuel Llinás (The Pennsylvania State University, USA) and Alfred Cortes (ISGLobal, Spain) for kindly share protocols, Verónica Sanz Serrano (IPBLN, Spain) and the Microscopy facility, Flow cytometry service and Genomic Unit at the IPBLN for technical support.

## Notes

### Competing Interest Statement

The authors have declared no competing interest.

## References

1. World malaria report 2025: addressing the threat of antimalarial drug resistance. Geneva: World Health Organization. https://www.who.int/teams/global-malaria-programme/reports/world-malaria-report-2025 (2025, accessed 4 August 2026).

2. Gardner MJ, Hall N, Fung E, White O, Berriman M, et al. Genome sequence of the human malaria parasite Plasmodium falciparum. Nature 2002;419:498–511.

3. Lee MCS, Lindner SE, Lopez-Rubio J-J, Llinás M. Cutting back malaria: CRISPR/Cas9 genome editing of Plasmodium. Brief Funct Genomics 2019;18:281–289.

4. Zhang M, Wang C, Otto TD, Oberstaller J, Liao X, et al. Uncovering the essential genes of the human malaria parasite Plasmodium falciparum by saturation mutagenesis. Science 2018;360:eaap7847.

5. Kudyba HM, Cobb DW, Vega-Rodríguez J, Muralidharan V. Some conditions apply: Systems for studying Plasmodium falciparum protein function. PLOS Pathog 2021;17:e1009442.

6. Baum J, Papenfuss AT, Mair GR, Janse CJ, Vlachou D, et al. Molecular genetics and comparative genomics reveal RNAi is not functional in malaria parasites. Nucleic Acids Res 2009;37:3788–3798.

7. Mueller A-K, Hammerschmidt-Kamper C, Kaiser A. RNAi in Plasmodium. Curr Pharm Des;20:278–283.

8. Hentzschel F, Mitesser V, Fraschka SA-K, Krzikalla D, Carrillo EH, et al. Gene knockdown in malaria parasites via non-canonical RNAi. Nucleic Acids Res 2020;48:e2.

9. Armstrong CM, Goldberg DE. An FKBP destabilization domain modulates protein levels in Plasmodium falciparum. Nat Methods 2007;4:1007–1009.

10. Ganesan SM, Falla A, Goldfless SJ, Nasamu AS, Niles JC. Synthetic RNA–protein modules integrated with native translation mechanisms to control gene expression in malaria parasites. Nat Commun 2016;7:10727.

11. Prommana P, Uthaipibull C, Wongsombat C, Kamchonwongpaisan S, Yuthavong Y, et al. Inducible Knockdown of Plasmodium Gene Expression Using the glmS Ribozyme. PLOS ONE 2013;8:e73783.

12. Kreidenweiss A, Hopkins AV, Mordmüller B. 2A and the Auxin-Based Degron System Facilitate Control of Protein Levels in Plasmodium falciparum. PLOS ONE 2013;8:e78661.

13. Birnbaum J, Flemming S, Reichard N, Soares AB, Mesén-Ramírez P, et al. A genetic system to study Plasmodium falciparum protein function. Nat Methods 2017;14:450–456.

14. Collins CR, Das S, Wong EH, Andenmatten N, Stallmach R, et al. Robust inducible Cre recombinase activity in the human malaria parasite Plasmodium falciparum enables efficient gene deletion within a single asexual erythrocytic growth cycle. Mol Microbiol 2013;88:687–701.

15. Jones ML, Das S, Belda H, Collins CR, Blackman MJ, et al. A versatile strategy for rapid conditional genome engineering using loxP sites in a small synthetic intron in Plasmodium falciparum. Sci Rep 2016;6:21800.

16. Ramaprasad A, Blackman MJ. A scaleable inducible knockout system for studying essential gene function in the malaria parasite. Nucleic Acids Res 2025;53:gkae1274.

17. Konermann S, Lotfy P, Brideau NJ, Oki J, Shokhirev MN, et al. Transcriptome Engineering with RNA-Targeting Type VI-D CRISPR Effectors. Cell 2018;173:665–676.e14.

18. Yan WX, Chong S, Zhang H, Makarova KS, Koonin EV, et al. Cas13d Is a Compact RNA-Targeting Type VI CRISPR Effector Positively Modulated by a WYL-Domain-Containing Accessory Protein. Mol Cell 2018;70:327–339.e5.

19. Kushawah G, Hernandez-Huertas L, Prado JA-N del, Martinez-Morales JR, DeVore ML, et al. CRISPR-Cas13d Induces Efficient mRNA Knockdown in Animal Embryos. Dev Cell 2020;54:805–817.e7.

20. Buchman AB, Brogan DJ, Sun R, Yang T, Hsu PD, et al. Programmable RNA Targeting Using CasRx in Flies. CRISPR J 2020;3:164–176.

21. Hou S, Bi W, Liu W, Nie H, Jiang J, et al. Beyond DNA editing: how Cas13 redefined programmable RNA manipulation and what still limits its therapeutic promise. Nucleic Acids Res 2026;54:gkag616.

22. Ai Y, Liang D, Wilusz JE. CRISPR/Cas13 effectors have differing extents of off-target effects that limit their utility in eukaryotic cells. Nucleic Acids Res 2022;50:e65.

23. Li Y, Xu J, Guo X, Li Z, Cao L, et al. The collateral activity of RfxCas13d can induce lethality in a RfxCas13d knock-in mouse model. Genome Biol 2023;24:20.

24. Dalla Benetta E, López-Denman AJ, Li H-H, Masri RA, Brogan DJ, et al. Engineered Antiviral Sensor Targets Infected Mosquitoes. CRISPR J 2023;6:543–556.

25. Tong H, Huang J, Xiao Q, He B, Dong X, et al. High-fidelity Cas13 variants for targeted RNA degradation with minimal collateral effects. Nat Biotechnol 2023;41:108–119.

26. Adjalley SH, Lee MCS, Fidock DA. A Method for Rapid Genetic Integration into Plasmodium falciparum Utilizing Mycobacteriophage Bxb1 Integrase. In: Braman J (editor). In Vitro Mutagenesis Protocols: Third Edition. Totowa, NJ: Humana Press. pp. 87–100.

27. Moreno-Sánchez I, Hernández-Huertas L, Nahón-Cano D, Martínez- García PM, Treichel AJ, et al. Enhanced RNA-targeting CRISPR-Cas technology in zebrafish. Nat Commun 2025;16:2591.

28. Wei J, Lotfy P, Faizi K, Baungaard S, Gibson E, et al. Deep learning and CRISPR-Cas13d ortholog discovery for optimized RNA targeting. Cell Syst 2023;14:1087–1102.e13.

29. Wessels H-H, Méndez-Mancilla A, Guo X, Legut M, Daniloski Z, et al. Massively parallel Cas13 screens reveal principles for guide RNA design. Nat Biotechnol 2020;38:722–727.

30. Portugaliza HP, Llorà-Batlle O, Rosanas-Urgell A, Cortés A. Reporter lines based on the gexp02 promoter enable early quantification of sexual conversion rates in the malaria parasite Plasmodium falciparum. Sci Rep 2019;9:14595.

31. Knuepfer E, Napiorkowska M, van Ooij C, Holder AA. Generating conditional gene knockouts in Plasmodium – a toolkit to produce stable DiCre recombinase-expressing parasite lines using CRISPR/Cas9. Sci Rep 2017;7:3881.

32. Kafsack BFC, Rovira-Graells N, Clark TG, Bancells C, Crowley VM, et al. A transcriptional switch underlies commitment to sexual development in malaria parasites. Nature 2014;507:248–252.

33. Poran A, Nötzel C, Aly O, Mencia-Trinchant N, Harris CT, et al. Single- cell RNA sequencing reveals a signature of sexual commitment in malaria parasites. Nature 2017;551:95–99.

34. Bancells C, Llorà-Batlle O, Poran A, Nötzel C, Rovira-Graells N, et al. Revisiting the initial steps of sexual development in the malaria parasite Plasmodium falciparum. Nat Microbiol 2019;4:144–154.

35. Tibúrcio M, Yang ASP, Yahata K, Suárez-Cortés P, Belda H, et al. A Novel Tool for the Generation of Conditional Knockouts To Study Gene Function across the Plasmodium falciparum Life Cycle. mBio 2019;10:e01170–19.

36. de Azevedo MF, Gilson PR, Gabriel HB, Simões RF, Angrisano F, et al. Systematic analysis of FKBP inducible degradation domain tagging strategies for the human malaria parasite Plasmodium falciparum. PloS One 2012;7:e40981.

37. Muralidharan V, Oksman A, Iwamoto M, Wandless TJ, Goldberg DE. Asparagine repeat function in a Plasmodium falciparum protein assessed via a regulatable fluorescent affinity tag. Proc Natl Acad Sci 2011;108:4411–4416.

38. Nasamu AS, Falla A, Pasaje CFA, Wall BA, Wagner JC, et al. An integrated platform for genome engineering and gene expression perturbation in Plasmodium falciparum. Sci Rep 2021;11:342.

39. Rajaram K, Liu HB, Prigge ST. Redesigned TetR-Aptamer System To Control Gene Expression in Plasmodium falciparum. mSphere 2020;5:10.1128/msphere.00457-20.

40. Le Roch KG, Zhou Y, Blair PL, Grainger M, Moch JK, et al. Discovery of gene function by expression profiling of the malaria parasite life cycle. Science 2003;301:1503–1508.

41. Bozdech Z, Llinás M, Pulliam BL, Wong ED, Zhu J, et al. The Transcriptome of the Intraerythrocytic Developmental Cycle of Plasmodium falciparum. PLOS Biol 2003;1:e5.

42. Serrano-Durán R, López-Farfán D, Gómez-Díaz E. Epigenetic and Epitranscriptomic Gene Regulation in Plasmodium falciparum and How We Can Use It against Malaria. Genes 2022;13:1734.

43. Sinha A, Hughes KR, Modrzynska KK, Otto TD, Pfander C, et al. A cascade of DNA-binding proteins for sexual commitment and development in Plasmodium. Nature 2014;507:253–257.

44. Polino AJ, Nasamu AS, Niles JC, Goldberg DE. Assessment of Biological Role and Insight into Druggability of the Plasmodium falciparum Protease Plasmepsin V. ACS Infect Dis 2020;6:738–746.

45. Hart SK, Müller S, Wessels H-H, Méndez-Mancilla A, Drabavicius G, et al. Precise RNA targeting with CRISPR–Cas13d. Nat Biotechnol 2026;44:64–69.

46. Dijk MR van, Schaijk BCL van, Khan SM, Dooren MW van, Ramesar J, et al. Three Members of the 6-cys Protein Family of Plasmodium Play a Role in Gamete Fertility. PLOS Pathog 2010;6:e1000853.

47. Molina-Cruz A, Garver LS, Alabaster A, Bangiolo L, Haile A, et al. The Human Malaria Parasite Pfs47 Gene Mediates Evasion of the Mosquito Immune System. Science 2013;340:984–987.

48. Canepa GE, Molina-Cruz A, Barillas-Mury C. Molecular Analysis of Pfs47- Mediated Plasmodium Evasion of Mosquito Immunity. PLOS ONE 2016;11:e0168279.

