## Supplementary Data for "Engineering a High-Fidelity CRISPR-Cas13d System for Targeted RNA Depletion in *Plasmodium falciparum*"

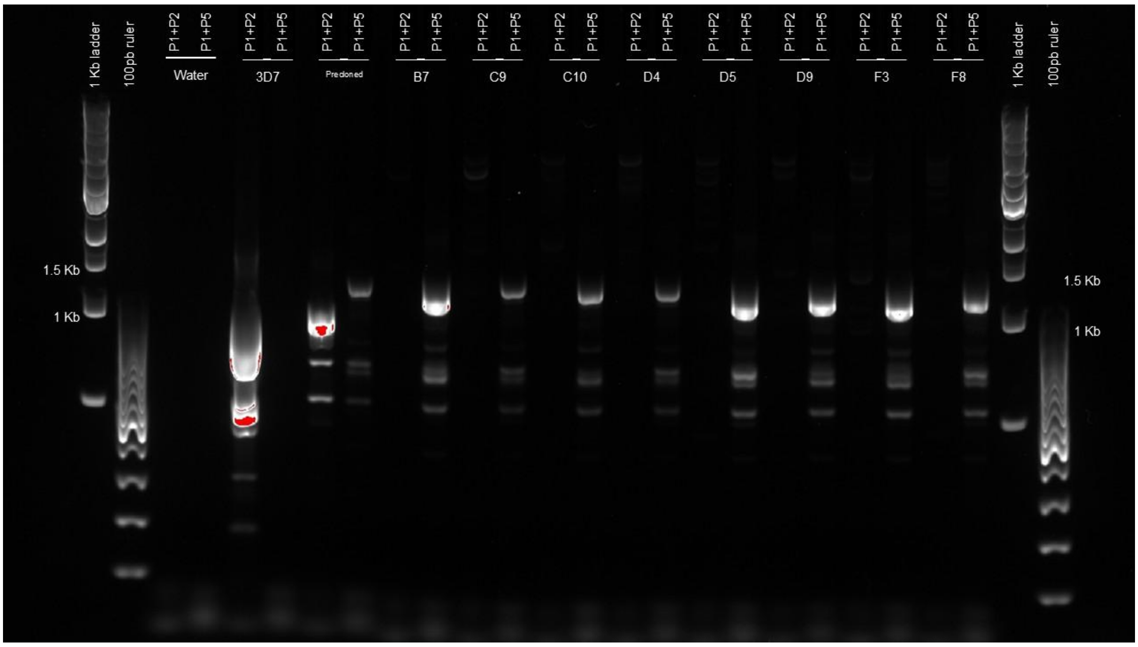


**Figure S1. Genotyping PCR validation of genetic modification in 3D7 HF-Cas13d clones.** Diagnostic PCR analysis confirming targeted integration of the HF-RfxCas13d expression cassette into the *pfs47* locus. Primer pair P1+P2 amplifies a 1,114 bp product corresponding to the wild-type *pfs47* locus. Primer pair P1+P5 amplifies a 1,257 bp fragment specific to the modified locus following successful integration. Water, 3D7 parental line, and pre-cloned transfectant parasite populations serve as negative, wild-type, and positive integration controls, respectively. Clones B7, C9, C10, D4 (hereinafter referred to as 3D7 HF-Cas13d), D5, D9, F3, and F8 denote individual parasite lines isolated by limiting dilution. Molecular weight markers (1 kb ladder and 100 bp ruler) serve as fragment length references.

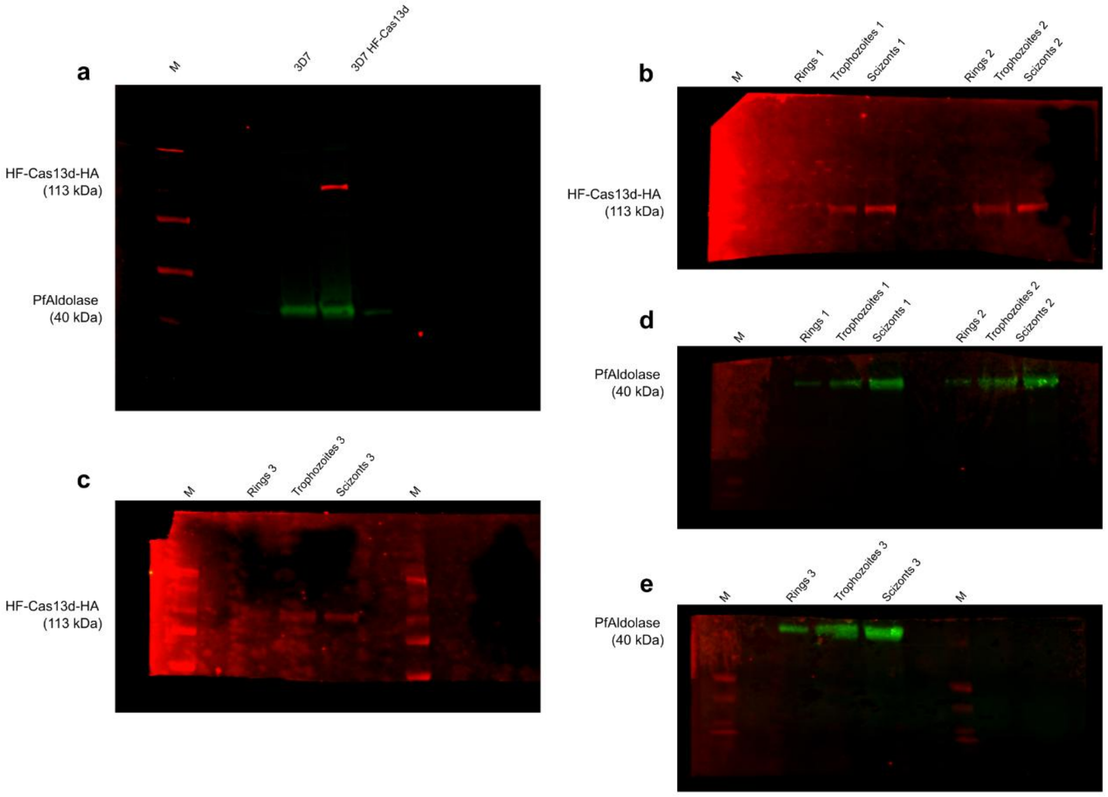


**Figure S2. Full-length western blot images detecting HF-RfxCas13d protein expression.** (**a**) Uncropped, full-length blot corresponding to HF-RfxCas13d detection in the 3D7 HF-Cas13d clonal line relative to the 3D7 parental line shown in Fig. 1c. (**b-c**) Uncropped, full-length blots representing independent biological replicates 1 (**b**), 2 (**b**), and 3 (**c**) used for densitometric quantification of HF-RfxCas13d protein expression across the IDC (rings, trophozoites, and schizonts) (Fig. 3h). (**d-e**) Uncropped, full-length blots representing independent biological replicates 1 (**d**), 2 (**d**), and 3 (**e**) showing PfAldolase loading control. Molecular weight marker (M).

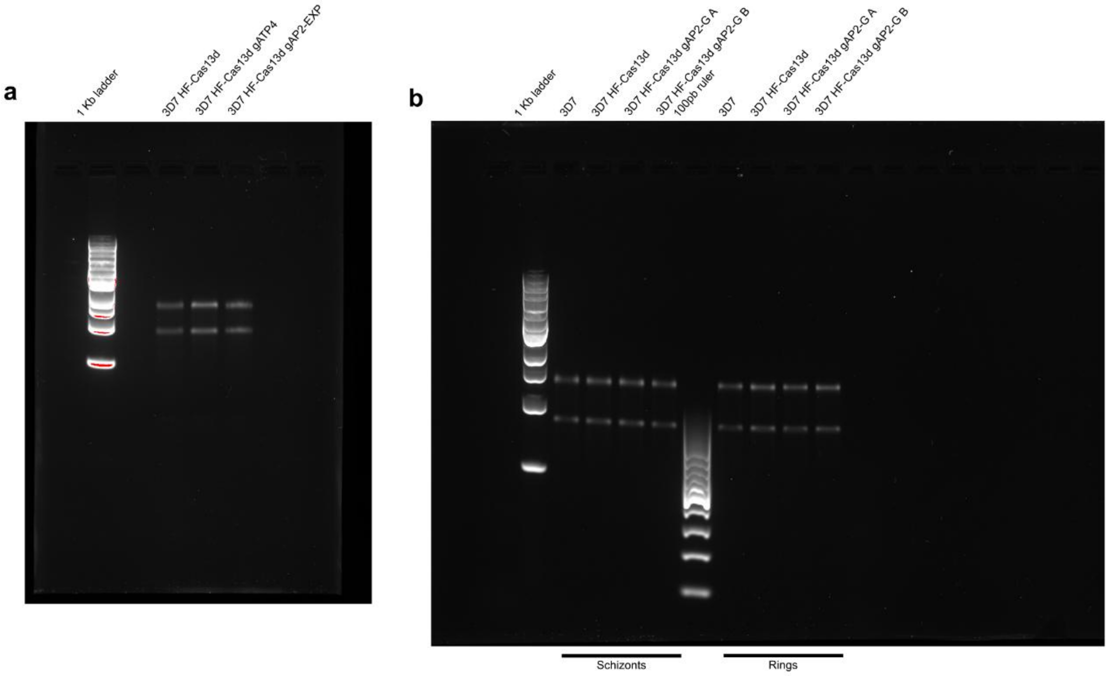


**Figure S3. Agarose gel electrophoresis for RNA integrity validation.** Verification of RNA integrity and absence of collateral ribosomal RNA degradation following HF-RfxCas13d activation. (**a**) Total RNA extracted from 3D7 HF-Cas13d parasites expressing gATP4 or gAP2-EXP. (**b**) Total RNA extracted from schizont (left) and ring (right) developmental stages of 3D7 HF-Cas13d parasites expressing gAP2-G A or gAP2-G B. 28S and 18S ribosomal RNA bands indicate the absence of non-specific transcript processing or non-specific bystander RNA cleavage in all conditions. Molecular weight markers 100 bp ruler and 1 kb ladder.

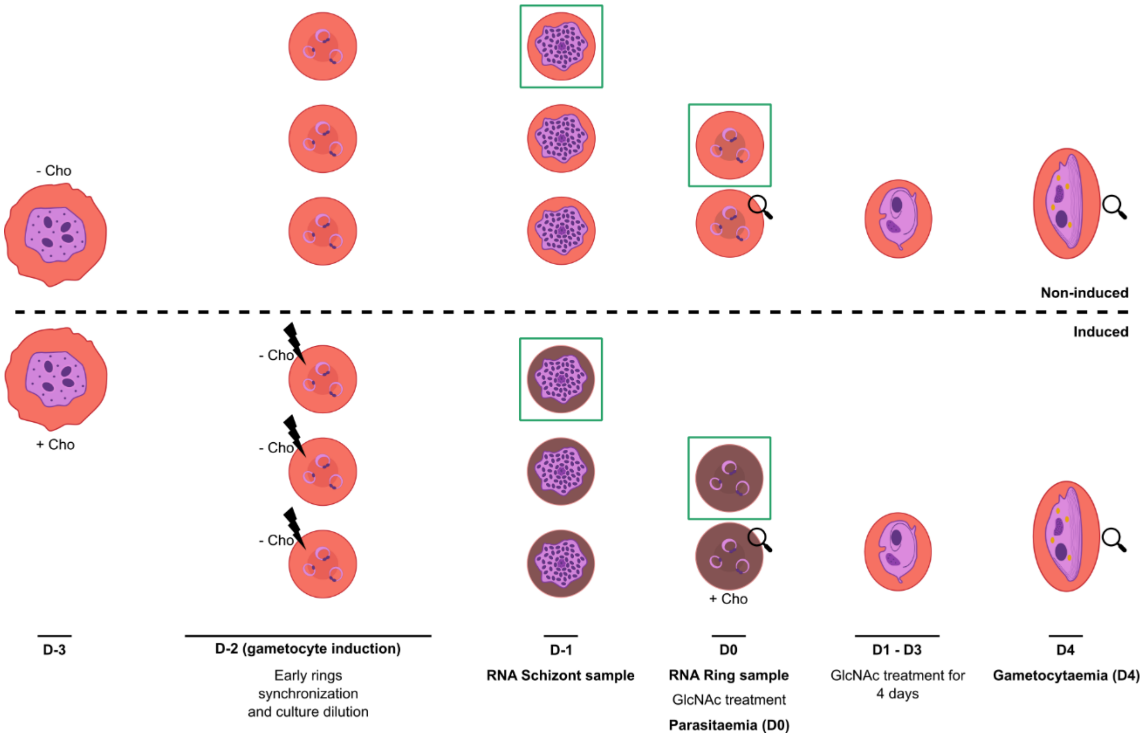


**Figure S4. Choline induction of sexual conversion.** Schematic representation of the cultures used for sexual induction assays. Non-induced parasites are shown at the top and induced parasites at the bottom. A lightning represents the induction by choline depletion, the green rectangles show the RNA sample uptake and the magnifying glasses illustrate the thin-smears preparation for parasitaemia and gametocytaemia quantifications.

**Table S1. Primers used in the study.**

| Primer name | Alternative name | Sequence | Annealing temperatures (ºC) |
|---|---|---|---|
| **P1** | *Pfs47ext-F* | ATTGCATACACATAAATATTTGTGTTGTAC | 57,3 / 56,5 |
| **P2** | *Pfs47int-R* | CACATACGTATTGTGTTGAGCTTAATAG | 57,3 |
| **P3** | *Pfs47int-F* | CTAATGTTAAGCCAACTGTAGTTGGG | 57,3 |
| **P4** | *Pfs47ext-R* | GATGCGATATGTAATTCCATTACTGC | 57,3 / 53,5 |
| **P5** | *cam3'-R* | CCATTTACACATAAATGTCACACAAAAGAG | 56,5 |
| **P6** | *Bip5'-F* | CAACAATTTTCTTAAAATTTGTACG | 53,5 |
| **P7** | *atp4-A-F* | TGAAAAAGATGAAGTGGTACCTG | 54 |
| **P8** | *atp4-A-R* | GCGACTCCTTCAACAACTTC | 54 |
| **P9** | *atp4-B-F* | ACTTTAAAATGGGGAATACATGG | 54 |
| **P10** | *atp4-B-R* | ACTATATCCTACTTTAGCAGCAG | 54 |
| **P11** | *exp-A-F* | GTTAGCCAAGAACCAGCAGC | 54 |
| **P12** | *exp-A-R* | CCACGCACACATACGCTTAT | 54 |
| **P13** | *exp-B-F* | CACCTGATCATAGTCTTAAAGGG | 54 |
| **P14** | *exp-B-R* | GTGATGAACAAGAACATGGC | 54 |
| **P15** | *ap2g-B-F* | TCTCATTCTTCAAATAGCCTTG | 54 |
| **P16** | *ap2g-B-R* | TCAGCATCGTCATCATTTAAC | 54 |
| **P17** | *ap2g-A-F* | AATGCAAACAATTTCAACAATGG | 54 |
| **P18** | *ap2g-A-R* | AGCTGTTGGGAACATATTCATAG | 54 |
| **P19** | *serr-F* | AAGTAGCAGGTCATCGTGGTT | 54 |
| **P20** | *serr-R* | TTCGGCACATTCTTCCATAA | 54 |
| **P21** | *uce-F* | GGTGTTAGTGGCTCACCAATAGGA | 54 |
| **P22** | *uce-R* | GTACCACCTTCCCATGGAGTA | 54 |
| **P23** | *Cas13d_F* | CATGGCGAGCGAGGCCAGCATC | 63 |
| **P24** | *Cas13d_PstI_R* | TCTTATCTGCAGGTTAAGCGTAATCTGGAACATC | 63 |
| **P25** | *Cas13d_F2* | GAAACTGCTGTGTGTGCCTTTC | - |
| **P26** | *Cas13d_R* | CCACGATCTTTTCCAGCCTG | - |
| **P27** | *DHFR_BamHI_F* | TAAGCAGGATCCATGGTTGGTTCGCTAAACTGC | 54,5 |
| **P28** | *DHFR_HindIII_R* | CGCCGCAAGCTTTTAATCATTCTTCTCATATACTTC | 54,5 |
| **P29** | *U6_pUF_Cpf1* | GACATTTGGATTTCTACACATC | - |
| **P30** | *targetRfx-R* | AGGTCTTCTCGAAGACCCG | - |

**Table S2. gRNA used in the study.**

| Gene Target | Gene ID | gRNA name | Spacer Length | Spacer Sequence | Forward Oligo | Reverse Oligo |
|---|---|---|---|---|---|---|
| ***atp4*** | PF3D7_1211900 | gATP4 A | 30 nt | AATGCAAGACTAGCTACAGCAGCAACTAAC | aaacAATGCAAGACTAGCTACAGCAGCAACTAAC | aaaaGTTAGTTGCTGCTGTAGCTAGTCTTGCATT |
| ***atp4*** | PF3D7_1211900 | gATP4 B | 23 nt | TATATCCTACTTTAGCAGCAGCA | aaacTATATCCTACTTTAGCAGCAGCA | aaaaTGCTGCTGCTAAAGTAGGATATA |
| ***ap2-exp*** | PF3D7_1466400 | gAP2-EXP A | 30 nt | ACTACTACTGTTTCAGCTGCTGGTTCTTGG | aaacACTACTACTGTTTCAGCTGCTGGTTCTTGG | aaaaCCAAGAACCAGCAGCTGAAACAGTAGTAGT |
| ***ap2-exp*** | PF3D7_1466400 | gAP2-EXP B | 23 nt | ATGTTGTTGTTCTAGGGGCATAC | aaacATGTTGTTGTTCTAGGGGCATAC | aaaaGTATGCCCCTAGAACAACAACAT |
| ***ap2-g*** | PF3D7_1222600 | gAP2-G A | 30 nt | AGGATTGTGATTATGTGCCATACCATTGTT | aaacAGGATTGTGATTATGTGCCATACCATTGTT | aaaaAACAATGGTATGGCACATAATCACAATCCT |
| ***ap2-g*** | PF3D7_1222600 | gAP2-G B | 23 nt | AGTTATAACATATGTGCGTTCCC | aaacAGTTATAACATATGTGCGTTCCC | aaaaGGGAACGCACATATGTTATAACT |

Note: Single-stranded oligonucleotides were synthesized with 4-nt 5′ overhangs (aaac on the forward strand, complementary to the BbsI-cleaved direct repeat DR30 site; aaaa on the reverse strand, complementary to the transcription terminator) to enable directional ligation into BbsI-digested pL6-DR30-mutDHFR
